# Optogenetic activation of the serotonergic neural circuit between the dorsal raphe nucleus and pre-Bötzinger complex contributes to inhibition of seizure-induced respiratory arrest in the DBA/1 mouse SUDEP model

**DOI:** 10.1101/2020.12.04.410969

**Authors:** HaiXiang Ma, Qian Yu, Yue Shen, XiTing Lian, LeYuan Gu, Han lu, HaiTing Zhao, Chang Zeng, Kazuki Nagayasu, HongHai Zhang

## Abstract

Sudden unexpected death in epilepsy (SUDEP) is the leading cause of death among epilepsy patients, occurring even more frequently in cases with anti-epileptic drug resistance. However, the underlying mechanism of SUDEP remains elusive. Our previous study demonstrated that enhancement of serotonin (5-HT) synthesis by intraperitoneal (IP) injection of 5-hydroxytryptophan (5-HTP) significantly reduced the incidence of seizure-induced respiratory arrest (S-IRA) in a DBA/1 mouse SUDEP model. Given that the 5-HT2A receptor (5-HT2AR) plays an important role in mediating the respiration system in the brain, we hypothesized that 5-HT2AR plays a key role in S-IRA and SUDEP. To test this hypothesis, we examined whether the decreased incidence of S-IRA evoked by either acoustic stimulation or pentylenetetrazole (PTZ) injection following 5-HTP administration will be blocked by treatment with ketanserin (KET), a selective antagonist of 5HT2AR, in the DBA/1 mouse SUDEP model. We observed that the reduction in S-IRA by 5-HTP was significantly reversed by IP or intracerebroventricular injection of KET. Considering the localization of 5-HT2AR in the pre-Bötzinger complex (PBC), which plays a key role in regulating respiratory rhythm, we next examined whether KET acts on 5-HT2AR in the PBC. To test this hypothesis, we activated the neural circuit between the dorsal raphe nucleus (DR) and PBC using optogenetics technology. We observed that stimulation of TPH2-ChETA-expressing neurons in the DR reduced the incidence of S-IRA evoked by PTZ, and this suppressant effect was significantly reversed by administration of KET in the bilateral PBC with no changes in electroencephalogram activity. The neural circuit between the DR and PBC was confirmed by injection of cholera toxin subunit B555 (CTB-555), a nerve tracer, in the DR or PBC separately. Calcium signaling evoked by PTZ within neurons of the PBC during seizures was significantly reduced by photostimulation of the DR. Taken together, our findings suggest that 5-HT2AR plays a critical role in regulating S-IRA and targeting the serotonergic neural circuit between the DR and PBC is a promising approach to preventing SUDEP.

## 1. INTRODUCTION

Continued research has demonstrated that sudden unexpected death in epilepsy (SUDEP) is the leading cause of death among patients with epilepsy, occurring even more frequently among patients with antiepileptic drug resistance ^1–4^. Currently, respiratory dysfunction during seizures is regarded as a leading mechanism of SUDEP. The latest advancements in understanding the pathogenesis of SUDEP revealed that cardiopulmonary dysfunction plays an important role in the occurrence of SUDEP ^1–5^. Some studies have indicated that several selective serotonin (5-HT) reuptake inhibitors (SSRIs) can prevent S-IRA evoked by generalized audiogenic seizures (AGSz) in DBA/1 mice by elevating 5-HT levels in the synaptic cleft ^17–19^. However, due to the limitations of animal SUDEP models, the role of 5-HT synthesis and the effects of targeting 5-HT receptors in the brain in modulating S-IRA and SUDEP remain unclear.

It had been accepted that tryptophan hydroxylase-2 (TPH2) in brain can convert l-tryptophan to 5-hydroxytryptophan (5-HTP), which can be further converted to 5-HT by aromatic L-amino acid decarboxylase ^22–24^. A previous study showed that TPH2 is the rate-limiting enzyme in 5-HT synthesis in brain ^22^. Our group previously tested the correlation between TPH2 protein expression and activity and found that TPH2 protein expression varied consistently with TPH2 activity in the same model ^11^. Thus, the expression of TPH2 is vital for the conversion of endogenous 5-HTP to 5-HT, thereby reducing the incidence of S-IRA and preventing SUDEP. However, it remains unknown how 5-HT mediates S-IRA and SUDEP in our mouse model.

Our previous research showed that seizure-induced respiratory arrest (S-IRA) can lead to the occurrence of SUDEP when evoked by either acoustic stimulation or pentylenetetrazole (PTZ) injection in a DBA/1 murine model ^5–12^, and the incidence of S-IRA evoked by acoustic stimulation or PTZ treatment was significantly reduced by treatment with 5-HTP in two models of SUDEP ^5^. However, the key target within the brain that mediates the 5-HTP–induced reductions in S-IRA and SUDEP has yet to be identified. Considering evidence that 5-HT2A receptor (5-HT2AR) is a key factor mediating respiratory function in the brainstem ^14–15^, we questioned whether application of 5-HTP could reduce the incidence of S-IRA by targeting 5-HT2AR in the brain. If so, we hypothesized that the suppressive effects of 5-HTP against S-IRA could be reversed by ketanserin (KET), a selective antagonist for 5-HT2AR, offering 5-HT2AR a potential therapeutic target for preventing SUDEP.

To investigate the roles of 5-HTP and 5-HT2AR in the pathogenesis of S-IRA and SUDEP, we continued to apply acoustic stimulation and PTZ injection in the DBA/1 mouse SUDEP model to test the ability of KET to reverse the effects of 5-HTP observed in our previous study. Considering the localization of 5-HT2AR in the pre-Bötzinger complex (PBC), which plays a key role in regulating respiratory rhythm, we next examined whether KET acts on 5-HT2AR in the PBC ^12^. To test this hypothesis, we activated the neural circuit between the dorsal raphe nucleus (DR) and PBC using optogenetics technology. Based on our previous finding that S-IRA in DBA/1 mice can be significantly prevented by optogenetic activation of TPH2-Channelrhodopsin 2 (ChR2) neurons in the DR, we further activated TPH2-ChETA neurons in the DR of DBA/1 mice to observe whether the suppression of S-IRA in the PTZ injection model by photostimulation of the DR depends on the activation of 5-HT2AR located in the PBC. It turned out that the lower incidence of S-IRA evoked by PTZ by optogenetic activation of TPH2-ChETA neurons in the DR was remarkably reversed by injection of KET into the pre-Bötzinger complex. Meanwhile, the neural circuit between the DR and the pre-Bötzinger complex was confirmed by injection of tracer CTB-555 into the DR and the pre-Bötzinger complex, respectively. Thus, our findings suggested that activating the neural circuit between the DR and the PBC might contribute to preventing SUEP.

## 2. MATERIALS AND METHODS

### 2.1 Animals

All experimental procedures were in line with the National Institutes of Health Guidelines for the Care and Use of Laboratory Animals and approved by the Animal Advisory Committee of Zhejiang University. DBA/1 mice were housed and bred in the Animal Center of Zhejiang University School of Medicine and given rodent food and water ad libitum. For the acoustic stimulation murine model, DBA/1 mice were “primed” starting from postnatal days 26–28 to establish consistent susceptibility to audiogenic seizures and S-IRA. For the PTZ-evoked seizure model, PTZ was administered to non-primed DBA/1 mice at approximately 8 weeks of age. TPH2-ChETA (E123T mutation in Channelrhodopsin 2 [ChR2])–expressing mice were generated by viral delivery of pAAV-TPH2 PRO-ChETA-EYFP-WPRES-PAS under the control of the promoter of TPH2 into the DR, and TPH2-ChETA expression in the DR for 3 weeks and was confirmed by immunohistochemistry after finishing optogenetics experiments . The promoter for TPH2 that we used in the present study has been used previously to infect 5-HT neurons in the DR by our group to modulate the balance between reward and aversion ^13^.

### 2.2 Seizure induction and resuscitation

S-IRA was evoked by acoustic stimulation or intraperitoneal (IP) administration of PTZ, as previously described ^5–12^. For the acoustic stimulation model, each mouse was placed in a cylindrical plexiglass chamber in a sound-isolated room, and AGSz were evoked by an electric bell (96 dB SPL, Zhejiang People’s Electronics, China). Acoustic stimulation was given for a maximum duration of 60 s or until the onset of tonic seizures and S-IRA. For the PTZ-evoked seizure model, S-IRA was evoked in non-primed DBA/1 mice by IP administration of a single dose of PTZ (Cat #P6500; Sigma-Aldrich, St. Louis, MO) at a dose of 75 mg/kg. Mice with S-IRA were resuscitated within 5 s after the final respiratory gasp using a rodent respirator (YuYan Instruments, Shanghai, China).

### 2.3. Pharmacology experiment

#### 2.3.1 Effect of IP administration of KET on 5-HTP–mediated suppression of S-IRA evoked by acoustic stimulation

As shown in Fig 1A, susceptibility to S-IRA in primed DBA/1 mice was confirmed 24 h prior to treatment with 5-HTP (Cat #107751; Sigma-Aldrich) or vehicle. 5-HTP (200 mg/kg) or vehicle (saline) was administered IP once daily for 2 days, and induction of S-IRA was performed 90 min after the second administration. KET (Cat #8006; Sigma-Aldrich) at different doses or vehicle (25% dimethyl sulfoxide [DMSO]) was administered IP 30 min before acoustic stimulation. The effects of 5-HTP and KET on S-IRA were examined and digitally recorded for offline analysis.

**Figure 1.**
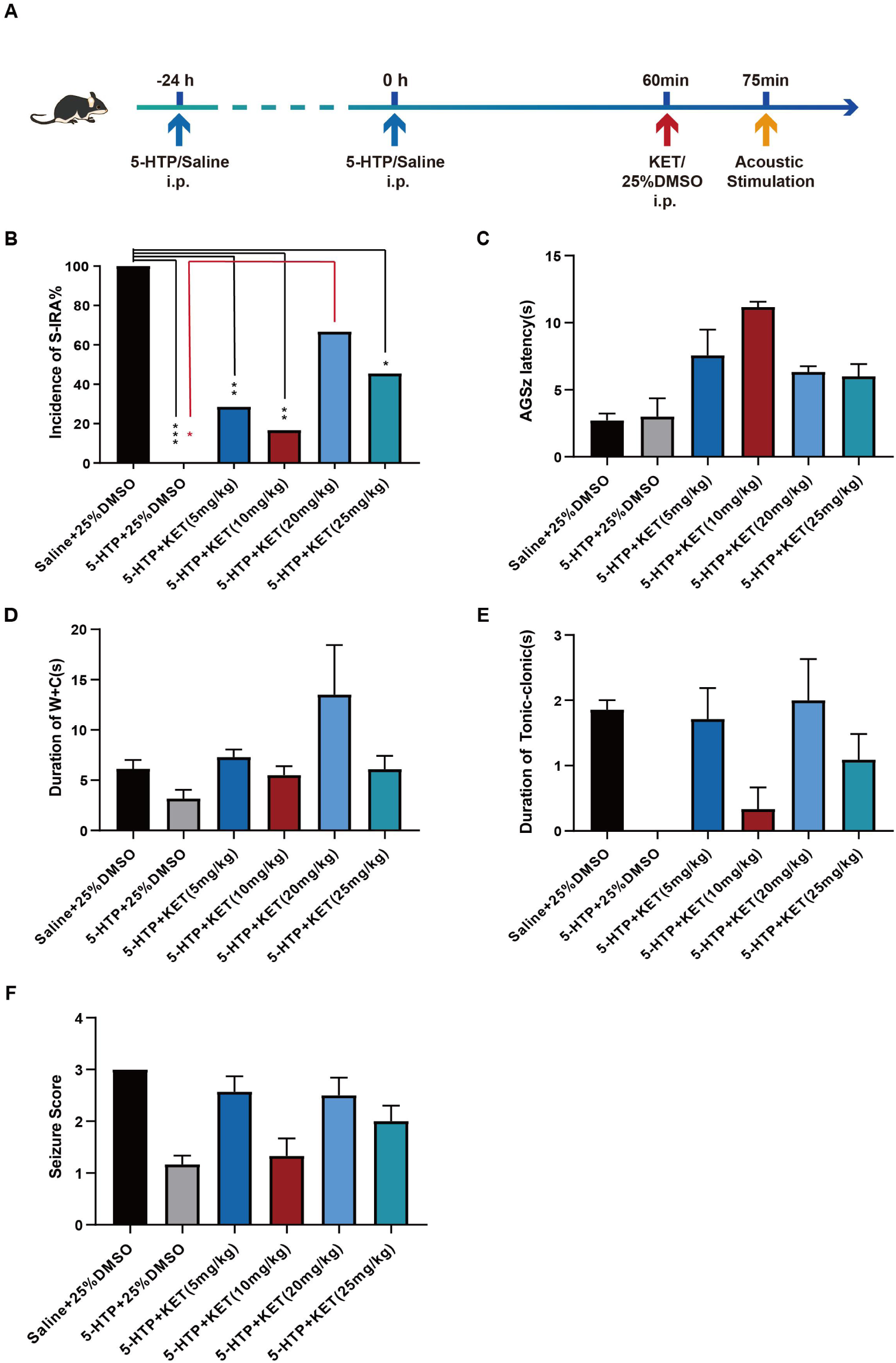
5-HTP-mediated reduction in S-IRA evoked by acoustic stimulation was significantly reversed by IP injection of KET. B. Compared with that in the group treated with saline and 25% DMSO, the incidence of S-IRA evoked by acoustic stimulation was markedly lower in groups treated with 5-HTP and 25% DMSO (n=7 and n=6, respectively; \**P*<0.05). Compared to that in the control group treated with vehicle, the incidence of S-IRA was significantly reduced in the group treated with 5-HTP and KET (5–10 mg/kg) (n=7 and n=6, respectively; *P*<0.01). However, no difference was found between the control group and the group treated with 5-HTP and KET (20 mg/kg) (n=7 and n=6, respectively; *P*>0.05). Furthermore, compared with that in the group treated with 5-HTP and 25% DMSO, the incidence of S-IRA in the group treated with 5-HTP and KET (20 mg/kg) was significantly increased (n=7 and n=6, respectively; \**P*<0.05). However, compared with that in the group treated with 5-HTP and 25% DMSO, the incidence of S-IRA in the group treated with 5-HTP and KET (25 mg/kg) was not significantly increased (n=7 and n=11, respectively; *P*>0.05). C-F, No intergroup differences were observed in latency to AGSz, duration of wild running and clonic seizures (W+C), duration of tonic-clonic seizures, and seizure scores (*P*>0.05).

#### 2.3.2 Effect of ICV administration of KET on 5-HTP–mediated suppression of S-IRA by PTZ

For intracerebroventricular (ICV) injection, an ICV guide cannula (O.D.I.41×I.D.O.0.25mm/M3.5, 62004, RWD Life Science Inc., China) was implanted into the right lateral ventricle (AP - 0.45 mm; ML - 1.0 mm; V - 2.50 mm) to enable microinjections as previously described ^9, 12^. KET or vehicle (25% DMSO) was administered by ICV injection 15 min prior to PTZ injection in DBA/1 mice in the same manner followed 5-HTP administration (Fig 2A). The incidence of S-IRA, latency to AGSz, duration of wild running and clonic seizures, duration of tonic-clonic seizures, and seizure scores were determined by offline analysis of video recordings ^5–10, 17–18^.The group treatments were as follows: 1) saline (IP) was administered 75 min prior to PTZ (75 mg/kg, IP) and 25% DMSO (2 μl, at a rate of 0.5 μl/min ICV) 15 min prior to PTZ injection as control; 2) 5-HTP (200 mg/kg, IP) was administered 75 min prior to PTZ (75 mg/kg, IP) and 25% DMSO (2 μl, at a rate of 0.5 μl/min ICV) 15 min prior to PTZ injection; and 3) 5-HTP (200 mg/kg, IP) was administered 75 min prior to PTZ (75 mg/kg, IP), with KET (9.15 nmol or 18.3 nmol, dissolved in 2 μl 25% DMSO, at a rate of 0.5 μl/min ICV) administered 15 min prior to PTZ injection.

**Figure 2.**
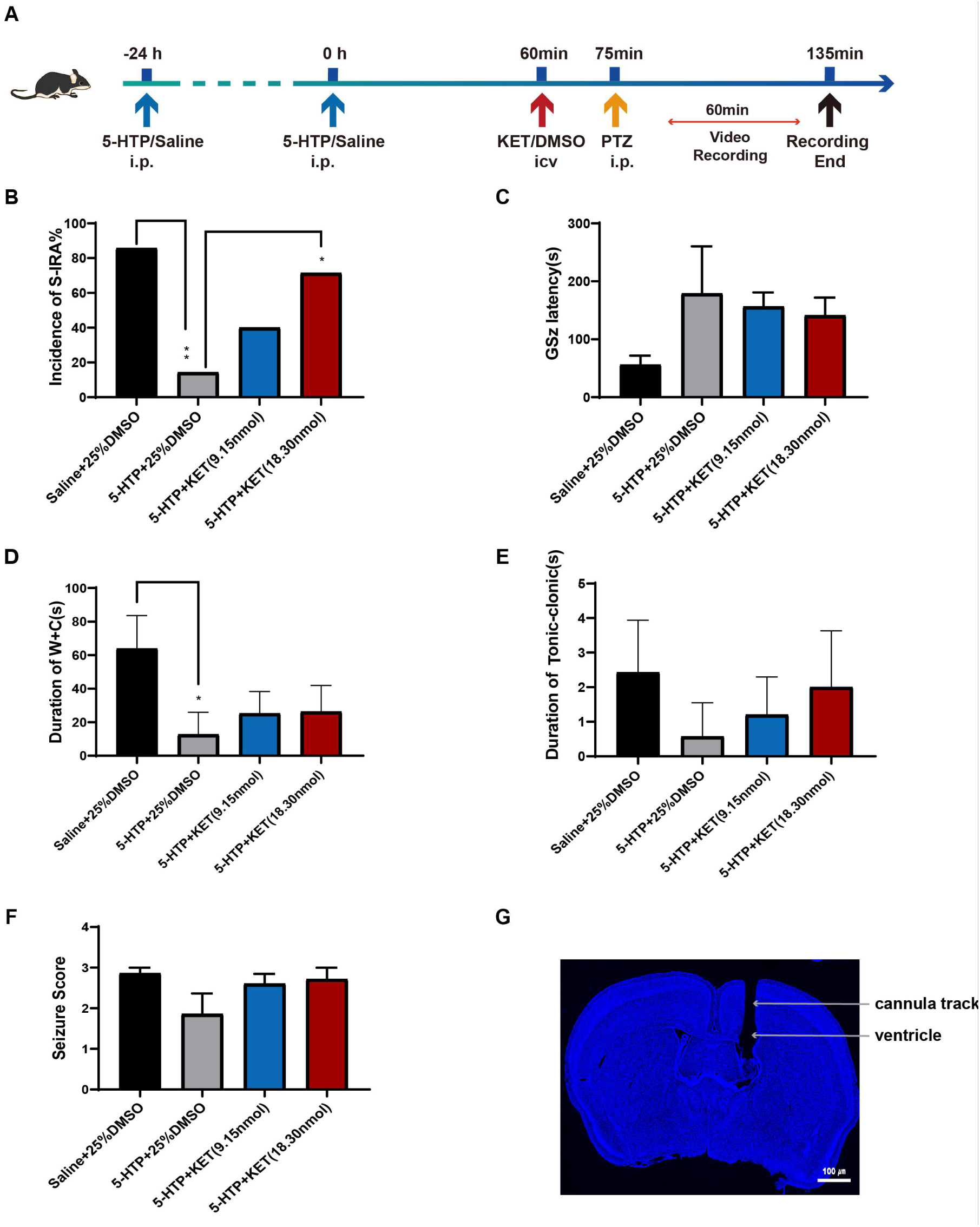
5-HTP–mediated reduction in PTZ-induced S-IRA was significantly reversed by ICV injection of KET. A, Compared to that in the control group, the incidence of PTZ-induced S-IRA was markedly lower in the group treated with 5-HTP and DMSO. However, no difference was observed between the control group and the groups treated with 5-HTP and KET (9.15 and 18.30 nmol). By contrast, the incidence of S-IRA was significantly reduced in the group treated with 5-HTP and ICV DMSO as compared with the groups treated with 5-HTP and KET (9.15 and 18.30 nmol). B–F, No intergroup differences were observed in latency to AGSz, duration of wild running plus clonic seizures (W+C), duration of tonic-clonic seizures, and seizure scores (*P*>0.05). G, Representative images of cannula implantation in a DBA/1 mice for ICV delivery. S-IRA, seizure-induced respiratory arrest; AGSz, audiogenic seizures; IP, intraperitoneal; DMSO, dimethyl sulfoxide. Data are mean ± SEM.

### 2.4 Optogenetics experiments

#### 2.4.1 Stereotactic surgery

DBA/1 mice at 8 weeks of age were anesthetized with 3.5% chloral hydrate and head-fixed in a stereotaxic apparatus (68018, RWD Life Science Inc., Shenzhen, China), as previously described ^7^. Throughout the entire surgical process, the body temperature of anesthetized mice was kept constant at 37°C using a heating pad. If the DBA/1 mice showed pain in response to a paw pinch, an additional 10% of the initial dosage of sodium pentobarbital was given to guarantee a painless state. For optogenetic viral delivery of pAAV-TPH2 PRO-ChETA-EYFP-WPRES-PAS, microinjection (100 nl, 40 nl/min) was performed using the following stereotaxic coordinates for the DR (AP − 4.55 mm, ML − 0.44 mm, DV − 2.80 mm, 10°right) based on the mouse brain atlas. Viruses were delivered via a gauge needle for the specification of 10ul (cat# 60700010, Gao Ge, Co., Ltd, ShangHai, China) by an Ultra Micro Pump (160494 F10E, WPI) over a period of 10 min; the syringe was not removed until 15–20 min after the end of infusion to allow diffusion of the viruses. Then, the optical fiber (FOC-W-1.25-200-0.37-3.0, Inper, Hangzhou, China) was implanted above the area (AP − 4.55 mm, ML − 0.44 mm, DV − 2.80 mm, 10°right) for 0.05 mm (AP − 4.55 mm, ML − 0.44 m, DV − 2.75 mm, 10°right). For ICV surgery, ICV guide cannula implantation was completed as described for the pharmacology experiment and was implanted with a headstage for EEG in the same mice as previously described ^12^. For microinjection of KET in the bilateral PBC, guide cannulas (O.D.0.48×I.D.0.34 mm/M3.5,62033, RWD Life Science Inc.) were implanted on both sides. CTB-555 (100 nl, 1 μg/μL, BrainVTA Technology Co. Ltd, Wuhan, China) was injected in the DR (AP − 4.55 mm, ML − 0.44 mm, DV − 2.80 mm, 10°right) or the right side of the PBC (AP − 6.80 mm, ML − 1.25 mm, DV − 4.95 mm), and we waited approximately 2 weeks for retrograde labeling of projection neurons. For the photostimulation of the bilateral PBC, pAAV-TPH2 PRO-ChETA-EYFP-WPRES-PAS was delivered in the bilateral PBC (AP − 6.80 mm, ML − 1.25 mm, DV − 4.95 mm) based on the mouse brain atlas for 3 weeks, and an optical fiber within a guide cannula (O.D.0.48×I.D.0.34mm/M3.5, 62033, RWD Life and FOC-W-1.25-200-0.37-3.0, Inper, Hangzhou, China) over 0.05 mm was implanted. For the photometry recordings, viral delivery of AAV2/9-mCaMKIIa-GCaMP6f-WPRE-pA via microinjection (100 nl,40 nl/min) was performed using the following stereotaxic coordinates for the bilateral PBC (AP − 6.80 mm, ML − 1.25 mm, DV − 4.95 mm) based on the mouse brain atlas for 3 weeks, and an optical fiber (FOC-W-1.25-200-0.37-3.0, Inper) over 0.05 mm was implanted.

#### 2.4.2 **P**hotostimulating of the DR-mediated suppression of S-IRA by PTZ and on EEG changes

pAAV-TPH2 PRO-ChETA-EYFP-WPRES-PAS was delivered into the DR of DBA/1 mice, and then an ICV guide cannula with a headstage for EEG was implanted for use for 3 weeks. The DBA/1 mice were divided into three groups. For the control group without photostimulation of the DR (n=7), ICV injection of vehicle (25% DMSO, 2 μl) was given in a uniform manner at the manner (0.5 μl/min) 30 min prior to IP injection of PTZ (75 mg/kg). For the group treated with photostimulation of the DR without ICV delivery of KET (n=6), ICV injection of the same concentration and volume of vehicle was given 15 min prior to photostimulation and 30 min prior to IP injection of PTZ (75 mg/kg). For another group treated with photostimulation of the DR and ICV delivery of KET (n=7), ICV injection of KET (total dose, 18.3 nmol) was given 15 min prior to photostimulation and 30 min prior to IP injection of PTZ (75 mg/kg). The incidence of S-IRA in each group was analyzed statistically. EEG recordings in the three groups were started 5 min before PTZ injection and ended 30 min after PTZ injection. EEG activity was statistically analyzed as previously described ^12^. The parameters for photostimulation of the DR were: blue-light, 465 nm, 20 Hz, 20-ms pulse width, 15 mW, and 20 min) was delivered by the laser (B12124, Inper) through a 200-μm optic fiber.

#### 2.4.3 Photometric analysis of DR-mediated suppression of S-IRA by PTZ after microinjection of KET in the bilateral PBC

pAAV-TPH2 PRO-ChETA-EYFP-WPRES-PAS was delivered into the DR of DBA/1 mice, and then and with tguide cannulas were implanted in the bilateral PBC for use for 3 weeks. For the group treated with photostimulation of the DR without administration of KET (n=7), microinjection of the vehicle (25% DMSO, 2 μl) into the bilateral PBC was given in a uniform manner (0.5 μl/min) 25 min prior to IP injection of PTZ (75 mg/kg). For another group treated with photostimulation of the DR with administration of KET (n=7), microinjection of KET with 200 nl (containing 0.4 µg KET) into the every unilateral PBC was given 25 min prior to IP injection of PTZ (75 mg/kg). Both groups received laser stimulation 10 min after microinjection. KET (0.8 µg in 400 nl) was given per mouse in the bilateral PBC. The parameters for photostimulation of the DR was the same to the ICV experiments.

#### 2.4.4 Photometric analysis of the incidence of PTZ-induced S-IRA with TPH2-ChETA infection of neurons of the bilateral PBC and microinjection of KET in the bilateral PBC

DBA/1 mice were used 3 weeks after viral delivery of pAAV-TPH2 PRO-ChETA-EYFP-WPRES-PAS into the bilateral PBC. For the control group treated without photostimulation (n=6), microinjection of the vehicle (25% DMSO, 2 μl) into the every unilateral PBC was given in a uniform manner (0.5 μl/min) 25 min prior to IP injection of PTZ (75 mg/kg). For the treated group treated with photostimulation of the bilateral PBC (n=7), microinjection of the same concentration and volume of vehicle into the every unilateral PBC was given 25 min prior to IP injection of PTZ (75 mg/kg). For the group treated with KET and photostimulation of the bilateral PBC (n=3), microinjection of KET (0.4 µg, 200 nl) into the every unilateral PBC was given 25 min prior to IP injection of PTZ (75 mg/kg). The parameters for photostimulation of the bilateral PBC were: blue-light, 465 nm, 20 Hz, 20-ms pulse width, 15 mW, and 20 min.

#### 2.4.5 Photometric analysis of Ca2^+^ activity of neurons in the bilateral PBC with photo-stimulation of TPH2-ChETA neurons in the DR in PTZ-induced S-IRA model

DBA/1 mice were used 3 weeks after viral delivery of pAAV-TPH2 PRO-ChETA-EYFP-WPRES-PAS or AAV2/9-mCaMKIIa-GCaMP6f-WPRE-pA (100 nl at a rate of 40 nl/min) into the DR and into the bilateral PBC. For the control group treated with pAAV-TPH2 PRO-ChETA-EYFP-WPRES-PAS in the DR without photostimulation and AAV2/9-mCaMKIIa-GCaMP6f-WPRE-pA in the bilateral PBC (n=3), photometry recordings were started 5 min prior to IP injection of PTZ (75 mg/kg) and ended 60 min after PTZ injection or until the death of mice if within 60 min. For the group treated with pAAV-TPH2 PRO-ChETA-EYFP-WPRES-PAS in the DR and photostimulation of the DR as well as AAV2/9-mCaMKIIa-GCaMP6f-WPRE-pA in the bilateral PBC (n=3), photometry recordings were started 5 min prior to IP injection of PTZ (75 mg/kg) and ended 60 min after PTZ injection or until the death of mice if within 60 min. The fiber photometry system (Inper, C11946) used a 488-nm diode laser. We segmented the data based on individual trials of seizure duration and determined the value of fluorescence change (ΔF/F) by calculating (F − F0)/F0.

### 2.5 Immunohistochemistry and histology

The placement of the optical fiber cannula tip for ICV microinjection of KET within the bilateral PBC in each mouse was verified by histology. The PBC region was identified by neurokinin-1 receptor (NK1R) staining. DBA/1 mice were sacrificed and perfused with phosphate-buffered saline (PBS) containing 4% paraformaldehyde (PFA). After saturation in 30% sucrose (24h), each mouse brain was sectioned into 30-μm-thick coronal slices with a freezing microtome (CM30503, Leica Biosystems, Buffalo Grove, IL, USA). The sections were first washed in PBS three times for 5 min each and then incubated in blocking solution containing 10% normal donkey serum (017-000-121, Jackson ImmunoResearch, West Grove, PA, USA), 1% bovine serum albumen (A2153, Sigma-Aldrich), and 0.3% Triton X-100 in PBS for 1 h at room temperature. Then, for c-fos or TPH2 staining in the DR, sections were incubated at 4°C overnight in a solution of rabbit anti-c-fos primary antibody (1:1000 dilution, 2250T Rabbit mAb /74620 Mouse mAb, Cell Signaling Technology, Danvers, MA, USA) or mouse anti-TPH2 primary antibody (1:100 dilution, T0678, Sigma-Aldrich). The secondary antibodies used were donkey anti-mouse Alexa 488 (1:1000; A32766, Thermo Fisher Scientific, Waltham, MA, USA), donkey anti-mouse Alexa 546 (1:1000; A10036, Thermo Fisher Scientific), or goat anti-rabbit cy5 (1:1000; A10523, Thermo Fisher Scientific), and slices were incubated in secondary antibody solutions for 2 h at room temperature. Similarly, for neurokinin 1 receptor (NK1R) or SA-2A R (5-HT-2AR) staining in the PBC, sections were incubated in a solution of rabbit anti-NK1 (1:1000 dilution, SAB4502913, Sigma-Aldrich) or mouse anti-SA-2A R (5-HT2AR) (1:100 dilution, sc-166775, Santa Cruz Biotechnology, Santa Cruz, CA, USA) at 4°C overnight followed by donkey anti-rabbit Alexa 488 secondary antibody (1:1000; A32766, Thermo Fisher Scientific), donkey anti-mouse Alexa 546 (1:1000; A10036, Thermo Fisher Scientific), or goat anti-mouse cy5 (1:400; A10524, Thermo Fisher Scientific) for 2 h at room temperature. After washing with PBS three times for 15 min each, the sections were mounted onto glass slides and incubated in DAPI (1:4000; Cat#C1002; Beyotime Biotechnology; Shanghai, China) for 7 min at room temperature. Finally, the glass slides were sealed using an anti-fluorescence attenuating tablet. All images were taken with a Nikon A1 laser-scanning confocal microscope (Nikon, Tokyo, Japan). The numbers of immunopositive cells were counted and analyzed using ImageJ (NIH, Bethesda, MD, USA). Notably, data from mice in which the implantation placement was outside of the targeted brain structure were not used in our experiments. Positively stained cells co-expressing c-fos, ChETA and TPH2 were counted as previously described ^7^.

### 2.6 Viral vectors

pAAV-TPH2 PRO-ChETA-EYFP-WPRES-PAS (AVV2/8) (viral titer: 1×10^13^ vg/ml), was purchased from Sheng BO, Co., Ltd. (Shanghai, China), and the sequences of vectors were designed by Kazuki Nagayasu (Department of Molecular Pharmacology Graduate School of Pharmaceutical Sciences, Kyoto University). CTB-555 (1 μg/μL) was purchased from Brain VTA Technology Co., Ltd. (Wuhan, China). AAV2/9-mCaMKIIa-GCaMP6f-WPRE-pA (viral titer: 1×10^14^ vg/ml) were purchased from Taitool Bioscience Co., Ltd. (Shanghai, China)

### 2.7 Statistical analysis

All data are presented as the mean ± standard error of the mean (SEM). Statistical analyses were performed using SPSS 23 (SPSS Software Inc., Chicago, IL, USA). The incidence rates of S-IRA in different groups were compared using Wilcoxon signed rank test. Seizure scores, the latency to AGSz, the duration of wild running and clonic seizures, and duration of tonic-clonic seizures were evaluated using one-way analysis of variance (ANOVA). The Mann–Whitney U test or the Kruskal–Wallis H test. Analysis of covariance (ANCOVA) was used to compare the numbers of c-fos–positive cells in the DR of DBA/1 mice with and without photostimulation. Statistical significance was inferred if *P*<0.05.

## **3.** Results

### 3.1 5-HTP–mediated suppression of S-IRA evoked by acoustic stimulation was reversed by IP injection of KET

Compared with that in the vehicle group in primed DBA/1 mice, the incidence of S-IRA evoked by acoustic stimulation was significantly reduced after IP delivery of 5-HTP at a dosage of 200 mg/kg (*P*<0.001). Moreover, compared with that in the vehicle group, the incidence of S-IRA in the group pre-treated with 5-HTP (200 mg/kg, IP) and KET (5, 10, or 25 mg/kg, IP) was significantly decreased (*P*<0.01, *P*< 0.05), indicating that these dosages of KET did not significantly reverse the prohibitive effect of 5-HTP. However, the incidence of S-IRA showed no difference between the vehicle group and the group treated with 5-HTP (200 mg/kg, IP) and KET (20 mg/kg, IP; *P*>0.05) and was significantly lower in the group treated with 5-HTP + 25% DMSO than in the group treated with KET (20 mg/kg, IP; *P*<0.05), Thus, the suppressive effect by 5-HTP against S-IRA was markedly reversed by IP administration of 20 mg/kg KET (Figure 1). (Fig 1).

### 3.2 5-HTP–mediated suppression of PTZ-induced S-IRA was reversed by ICV delivery of KET

Compared with that in the vehicle control group, the incidence of PTZ-induced S-IRA was significantly reduced in the group that received ICV injection of 5-HTP and 25% DMSO (*P*<0.05). Also, the incidence of PTZ-induced S-IRA was not significantly less in the group treated with 5-HTP and KET (9.15 nmol, ICV) compare with the vehicle group (*P*>0.05). The incidence of PTZ-induced S-IRA was significantly greater in the group treated with 5-HTP and KET (9.15 nmol, ICV) than in the group treated with 5-HTP and 25% DMSO (ICV, *P*<0.05), which suggested that the suppressive effect of 5-HTP against S-IRA was significantly reversed by KET at the ICV dosage of 9.15 nmol. Furthermore, compared with that in the vehicle control group, the incidence of PTZ-induced S-IRA in the group treated with 5-HTP + KET (18.30 nmol, ICV) was not significantly reduced (P>0.05), and the incidence of PTZ-induced S-IRA in the group treated with 5-HTP and 25% DMSO (ICV) was significantly reduced compared with that in the group treated with 5-HTP and KET (18.30 nmol, ICV, *P*<0.05). Thus, the suppressive effect of S-IRA against 5-HTP could be significantly reversed by KET at an ICV dosage of 18.30 nmol. No significant intergroup differences were observed in latency to AGSz, duration of wild running and clonic seizures, duration of tonic-clonic seizures, and seizure scores (AGSz latency: *P*=0.763, F=6; duration of wild running and clonic seizures: *P*=0.14, F=6; duration of tonic-clonic seizures: *P*=0.119, F=6; seizure score: *P*=0.304, F=6; Figure 2, C,D.E,F). The seizure scores for two models from different groups did not differ significantly (*P*>0.05), and no obvious influence on seizure behavior was observed within the reversal effects of KET (Figure 2).

### 3.3 ChETA expression was induced in the membrane of 5-HT neurons in the DR of DBA/1 mice

The virus for expression of pAAV-TPH2 PRO-ChETA-EYFP-WPRES-PAS, under the control of the TPH2 promoter, was delivered into the DR of wild-type DBA/1 mice with the age of 50 days to express for 3 weeks to create TPH2-ChETA-expressing mice. We first examined the expression of ChETA and TPH2 in 5-HT neurons in the DR of DBA/1 mice using immunohistochemistry (n=3). The expression of GFP, a surrogate marker for ChETA, was predominantly localized in the membranes of the cell body and axons of 5-HT neurons of the DR. TPH2 was predominantly confined to the cytosol of 5-HT neurons in the DR of DBA/1 mice. The ratio for merged expression of ChETA and TPH2 was consistent with our previous finding that the co-expression of ChR2 and TPH2 is located in the membranes of 5-HT neurons in the DR of DBA/1 mice ^7^ (Figures 3 and 4).

**Figure 3.**
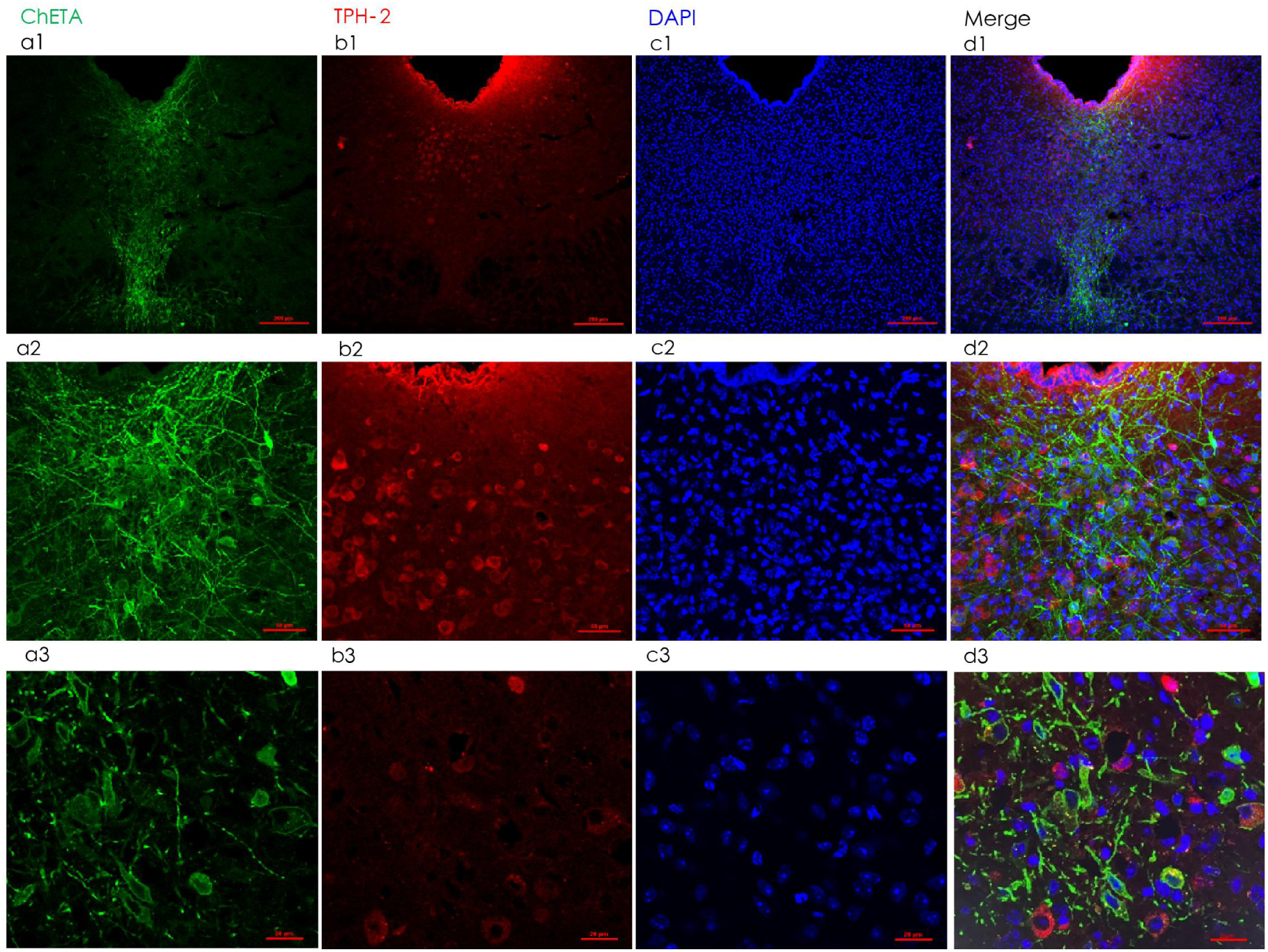
Localized expression of ChETA in 5-HT neurons in the DR of DBA/1 mice. a1, a2 and a3, Neuronal immunostaining of GFP, a surrogate marker for ChETA, in 5-HT neurons in the DR of a coronal brain slice. b1, b2 and b3, Immunostaining of TPH2, a key enzyme for 5-HT synthesis in the central nervous system. a1, b1, c3 and d1, merged images showing the co-expression of TPH2 and GFP in 5-HT neurons. These data demonstrate that ChETA was restrictively expressed on the surface of 5-HT neurons in the DR (n=3 mice). Confocal image magnifications: a1–d1, 10×; a2–d2, 20×; a3–d3, 40×.

**Figure 4.**
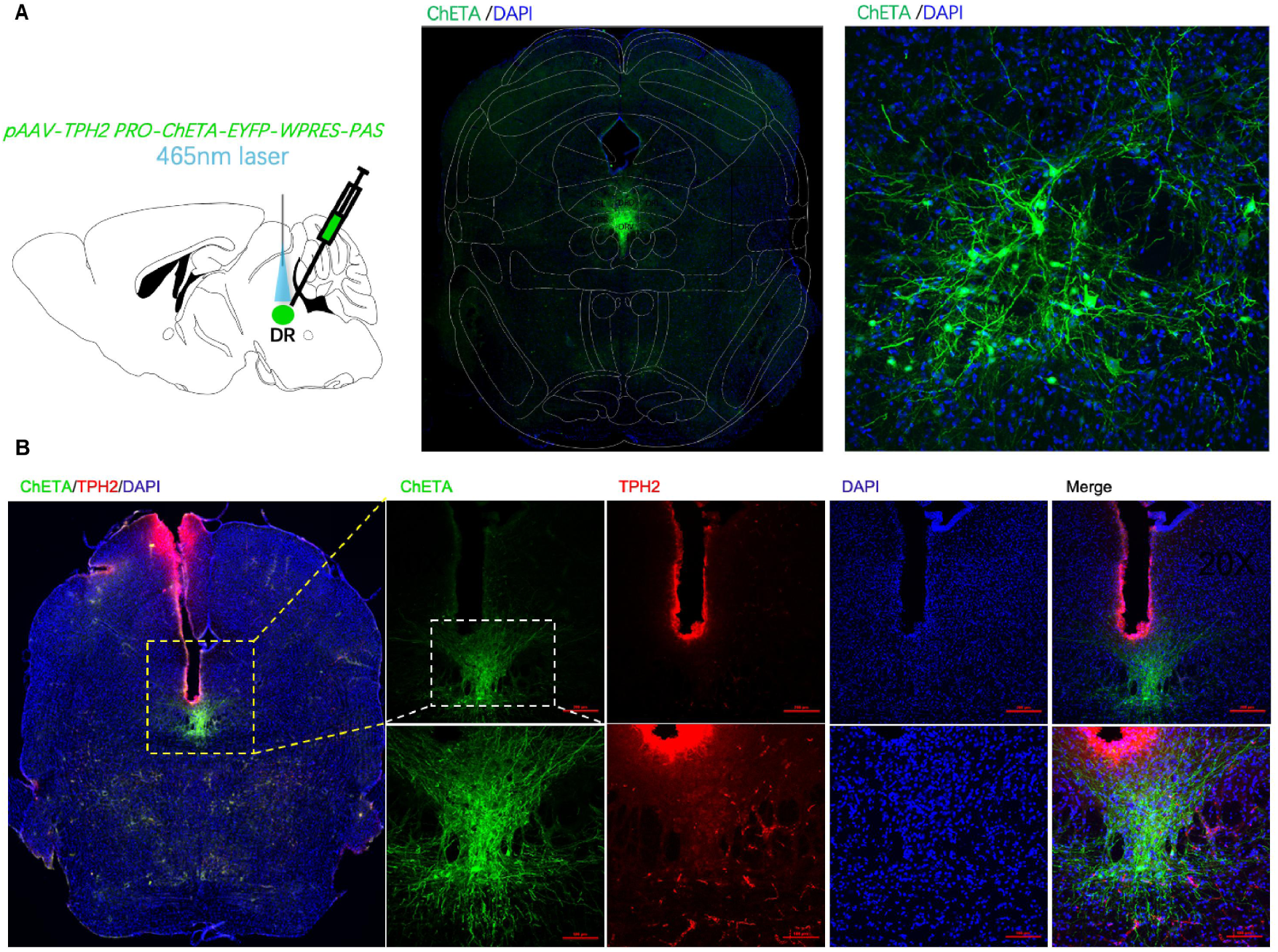
Placement of fiberoptic cannula tips in the DR. A, An example of a coronal brain slice, showing the location of an optic fiber cannula tip and the expression of ChETA in the DR of a DBA/1 mouse, according to the mouse atlas of Paxinos and Franklin (4th Edition, Paxinos and Franklin, 2013). B, Neuronal immunostaining of co-expression of ChETA and TPH2 in the DR of a DBA/1 mouse. No thermal injury due to photostimulation was observed in the area around the optic fiber tips.

### 3.4 Activation of 5-HT neurons in the DR reduced PTZ-induced S-IRA in DBA/1 mice and the lower incidence of S-IRA by photostimulating the DR this effect was significantly reversed by ICV delivery of KET

We examined the effect of selective enhancement of 5-HT neurotransmission on PTZ-induced S-IRA by applying photostimulation (blue light, 20-ms pulse duration, 20 Hz, 20 min) to 5-HT neurons in the DR of non-primed DBA/1 mice. Compared with the incidence of PTZ-induced S-IRA in DBA/1 mice without photostimulation (n=7), that after photostimulation of the DR for 20 min was significantly decreased (n=6, *P*<0.05). However, after ICV administration of KET, the incidence of PTZ-induced S-IRA was significantly increased under the same parameters of photostimulation of the DR. (n=7, *P*<0.05), indicating that the lower incidence of PTZ-induced S-IRA upon photostimulation of the DR was significantly reversed by blocking 5HT2AR receptor in the brain (Figure 5).

**Figure 5.**
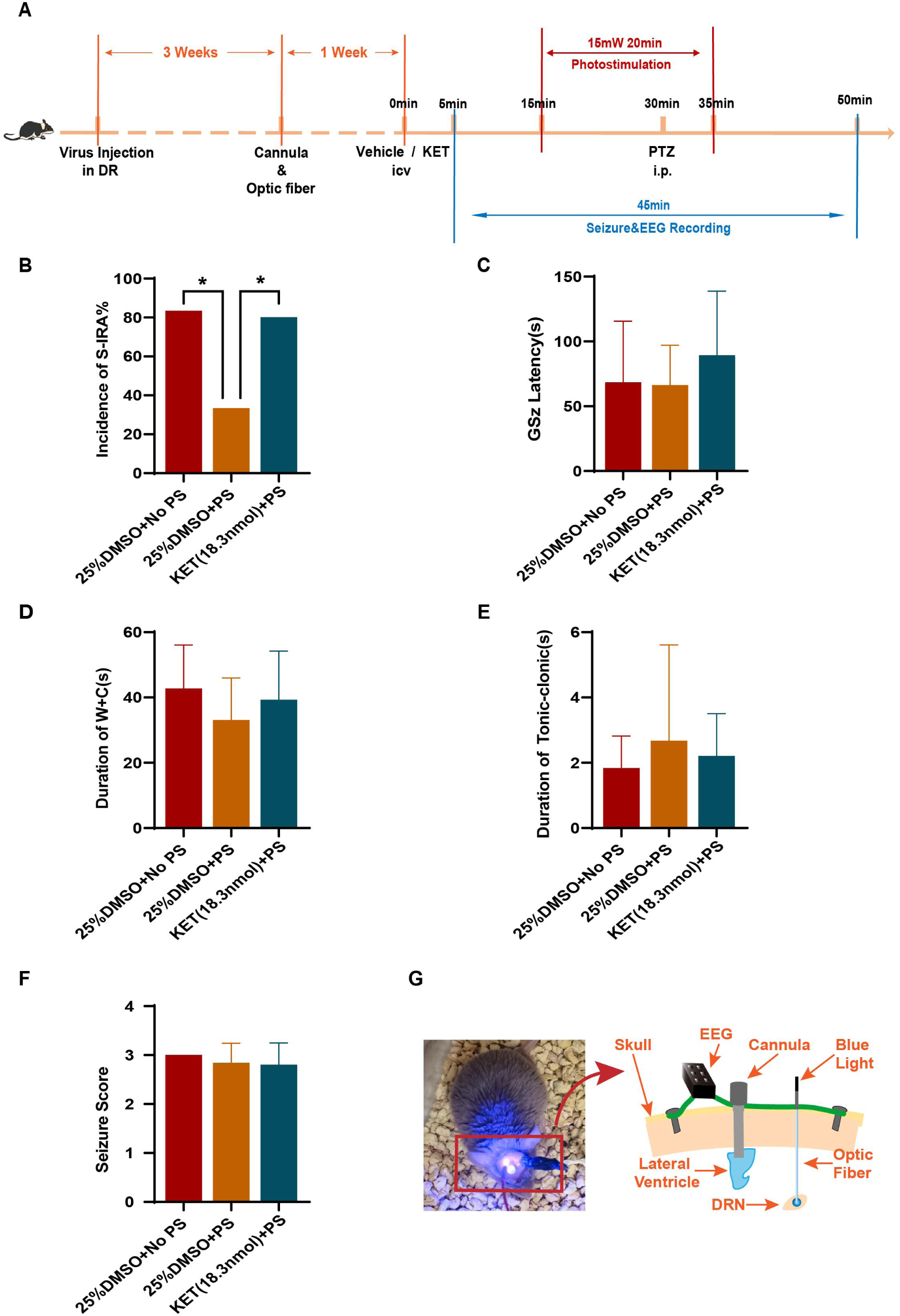
Optogenetic activation of TPH2-ChETA neurons in DR-mediated reduction of PTZ-induced S-IRA was significantly reversed by ICV injection of KET without changing seizure behavior. A, Compared with that in the control group treated with PTZ and no photostimulation, the incidence of PTZ-induced S-IRA was significantly reduced by photostimulation (n=7 and n=6, respectively; *P*<0.05). B, However, the lower incidence of S-IRA after photostimulation was remarkably reversed by ICV injection of KET at a dose of 18.3 nmol (n=6 and n=7, respectively; *P*<0.05). C-F, No obvious differences between groups were observed in the analysis of seizure score, duration of wild running and clonic seizure, AGSz latency, and duration of tonic seizures. G, Representative images of implanted EEG, ICV and optic fiber devices in a DBA/1 mouse. No PS = no photostimulation; PS = photostimulation.

### 3.5 Activation of 5-HT neurons in the DR produced differential effects on EEG activity in the S-IRA model with and without KET treatment

Based on the above findings in the same experimental groups, we further examined the effect of activating 5-HT neurons in the DR on EEG activity in our mouse model of S-IRA with and without KET treatment. Compared with that in the group not exposed to photostimulation, a EEG activity was significantly reduced upon photostimulation of 5-HT neurons in the DR of the mouse model. Analysis of the EEG wave data showed that the delta wave was significantly reduced by photostimulation, and this effect was reversed by KET treatment (P<0.05). No changes in the theta, alpha, beta, or gamma waves were apparent among the different treatment groups. These findings regarding EEG activity may reflect the specificity of 5-HT2AR in the brain for modulating S-IRA and SUDEP (Figure 6).

**Figure 6.**
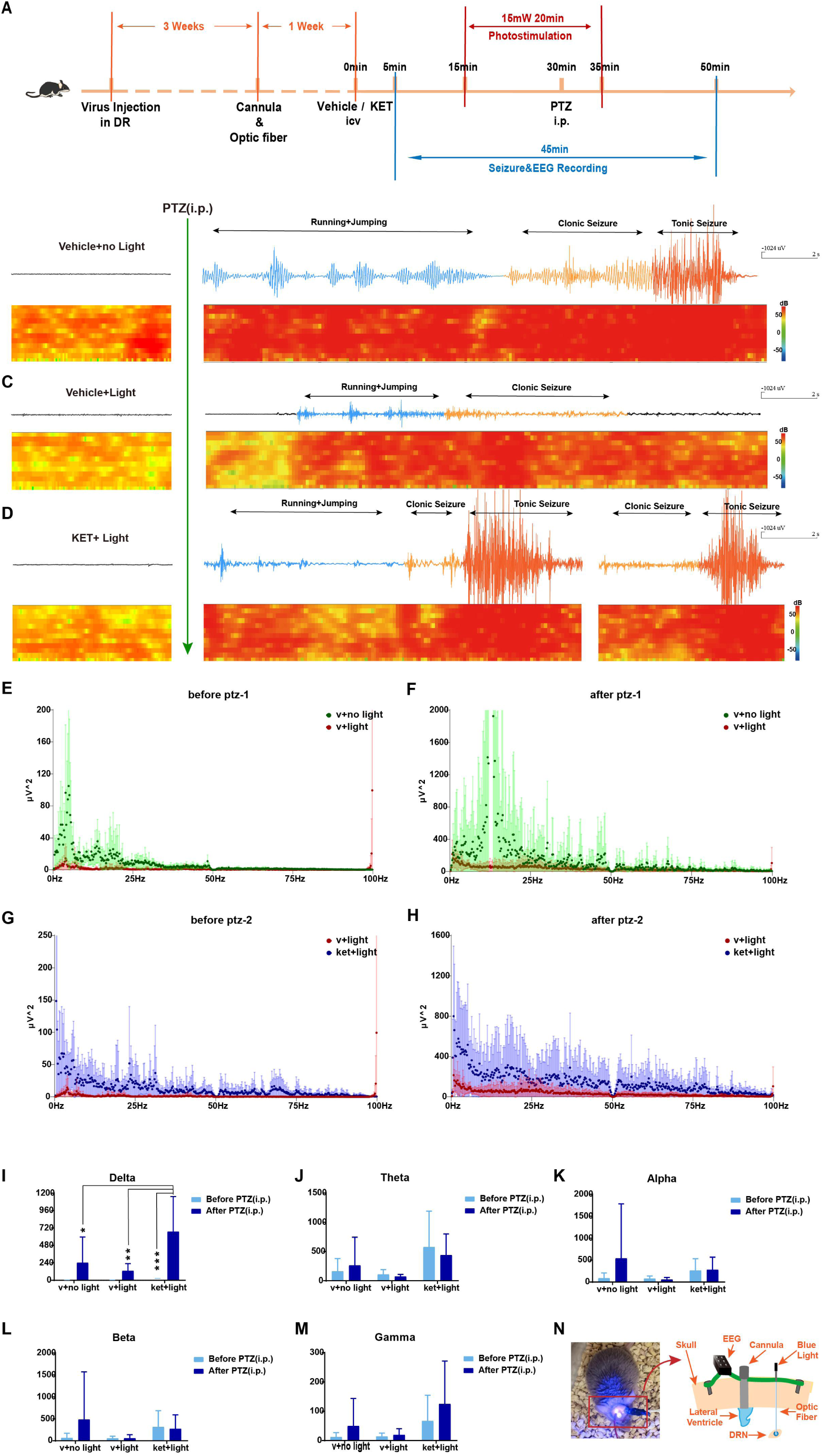
EEG activity upon optogenetic activation of TPH2-ChETA neurons in the DR-mediated reduction of PTZ-induced S-IRA and significant reversal of this reduction after ICV injection of KET. A. Schematic illustration of the pAAV-TPH2 PRO-ChETA-EYFP-WPRES-PAS being delivered into the DR of DBA/1 mice to implantion of optic fiber and cannula to receving the photostimulation of the DR and recoding the changing of EEG by ICV of KET. B-D Compared with that in the control group that received PTZ without photostimulation, the EEG activity during the clonic and tonic seizure stages was significantly suppressed in the group treated with PTZ and photostimulation. Furthermore, this suppression of EEG activity was significantly increased in the group treated with PTZ, photostimulation, and ICV microinjection of KET. E-M, Delta wave EEG activity was significantly reduced by light and reversed by KET. N, Representative images of implanted EEG, ICV, and optic fiber devices in a DBA/1 mouse. Light = photostimulation; No light = no photostimulation.

### 3.6 Reduction in S-IRA upon photostimulation of the DR was dependent on 5-HT2AR located in the PBC

Although the incidence of S-IRA can be reduced by 5-HTP and by photostimulation of the DR and this effect was significantly reversed by both IP and ICV injection of KET, it still was to determine whether the role of 5-HT2AR in the PBC in mediating the process of S-IRA and SUDEP remained unclear. The incidence of PTZ-induced S-IRA upon photostimulation of the DR was 14.28%, whereas that with photostimulation of the DR and microinjection of KET (400 nl) into the bilateral PBC was 85.71% (*P*<0.01). Subsequently, the existence of the neural circuit between the DR and the PBC was verified by using the nerve tracer CTB-555. Optogenetic activation of this neural circuit was shown to contribute to the inhibition of S-IRA, and 5-HT2AR in the PBC may be a specific target for preventing SUDEP (Figures 7–9). We speculated that the serotonergic neurotransmission resulting from optogenetic activation of the DR was accelerated between the DR and PBC, releasing the more content of 5-HT within PBC to activate 5-HT2AR to make abnormal respiratory rhythm recover the normal function against SIRA and SUDEP.

**Figure 7.**
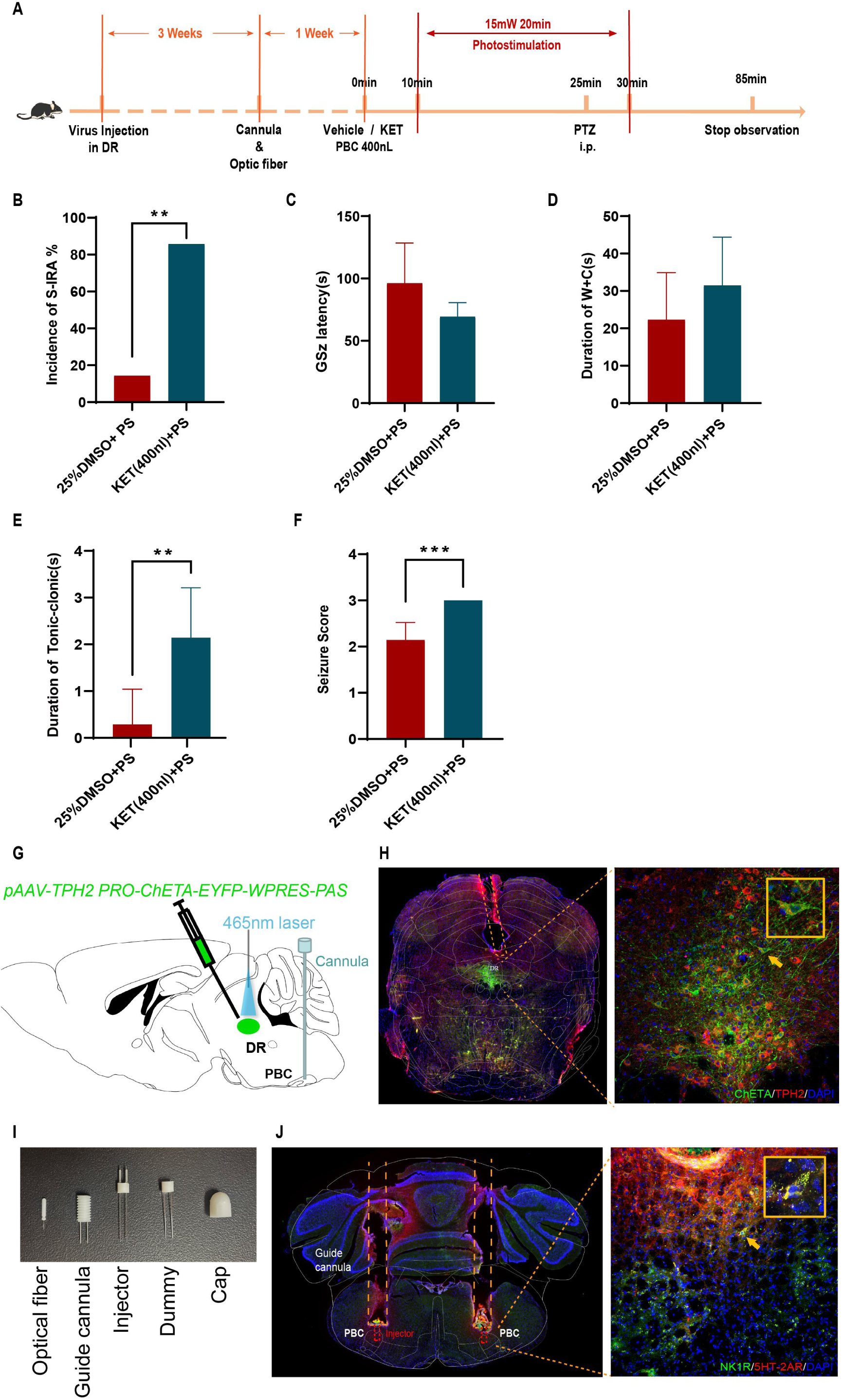
Optogenetic activation of TPH2-ChETA neurons in DR-mediated reduction of PTZ-induced S-IRA was significantly reversed by injection of KET into the PBC. A. Schematic illustration of the pAAV-TPH2 PRO-ChETA-EYFP-WPRES-PAS being delivered into the DR of DBA/1 mice to implantion of optic fiber and cannula to receving the photostimulation of the DR and oberving the changing of EEG by microinjection of KET in the PBC. B, Compared with that in the control group treated with PTZ and photostimulation, the incidence of PTZ-induce S-IRA was significantly reduced in the group that received photostimulation and microinjection of 400 nl KET into the bilateral PBC. C-D, No obvious differences were observed between treatment groups in the seizure score, duration of wild running and clonic seizure, AGSz latency, or duration of tonic seizures. E-F, Compared with that in the group treated with 25% DMSO + PS, the duration of tonic seizure and seizure score were significantly increased in the group treated with KET + PS (*P*<0.05 and *P*<0.01, respectively). No PS = no photostimulation; PS = photostimulation. G-J, Representative images of the delivery of pAAV-TPH2 PRO-ChETA-EYFP-WPRES-PAS into the DR and injection of KET into the bilateral PBC. H, Staining of ChETA, TPH2 and DAPI in the DR. I, All implantation devices in a DBA/1 mouse. J, The tracks for KET injection into the bilateral PBC and staining for NK1R, 5-HT2AR and DAPI in the bilateral PBC. No PS = no photostimulation; PS = photostimulation.

**Figure 8.**
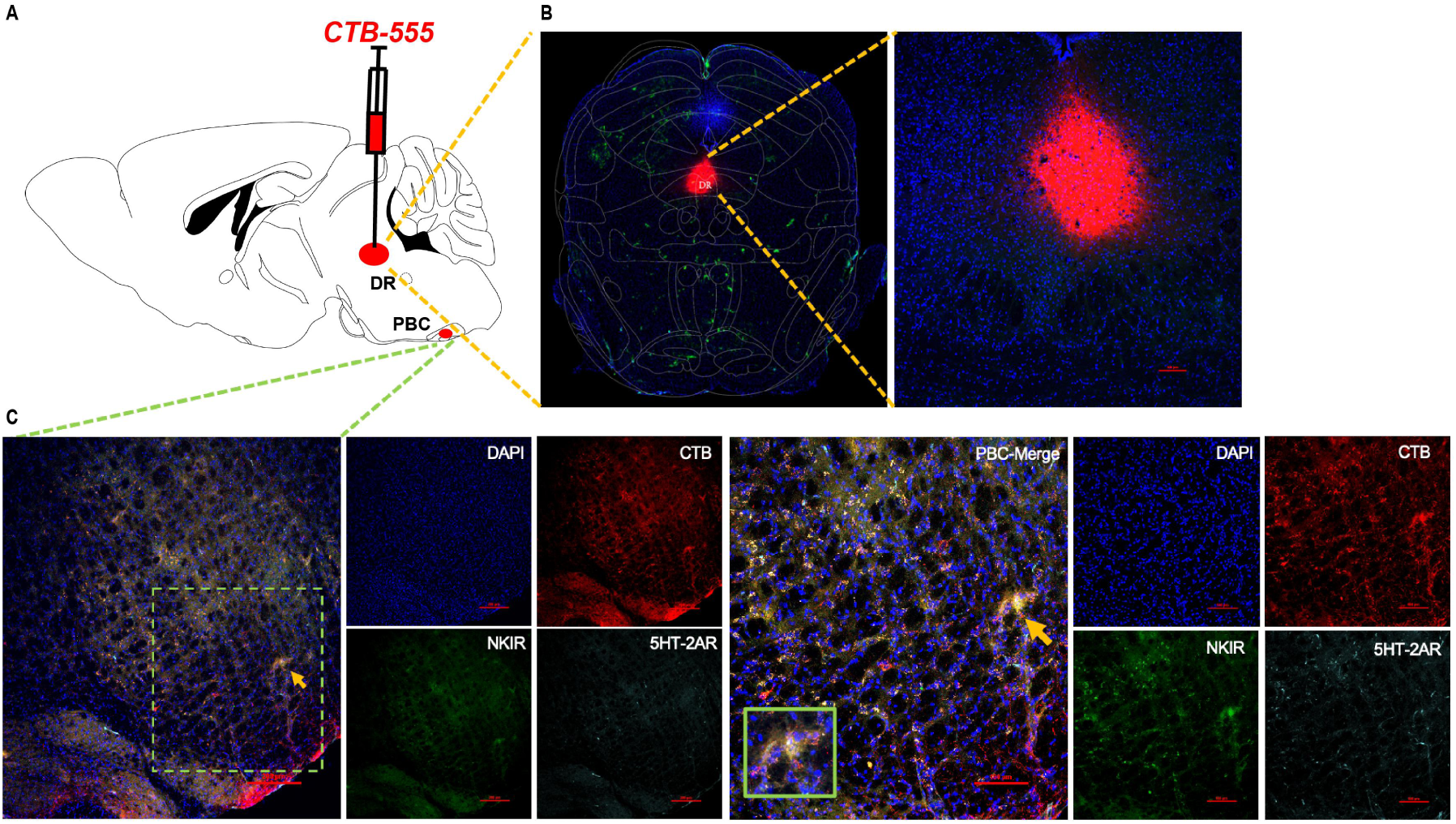
Neural projection from the DR to the PBC was established by application of the nerve tracer CTB555. A, Representative coronal brain slice, showing the location of CTB555 injection with the co-expression of TPH2 in the DR. B, Projection of the DR to the PBC with co-expression of CTB, NK1R, a marker for respiratory neurons, and 5HT2AR.

**Figure 9.**
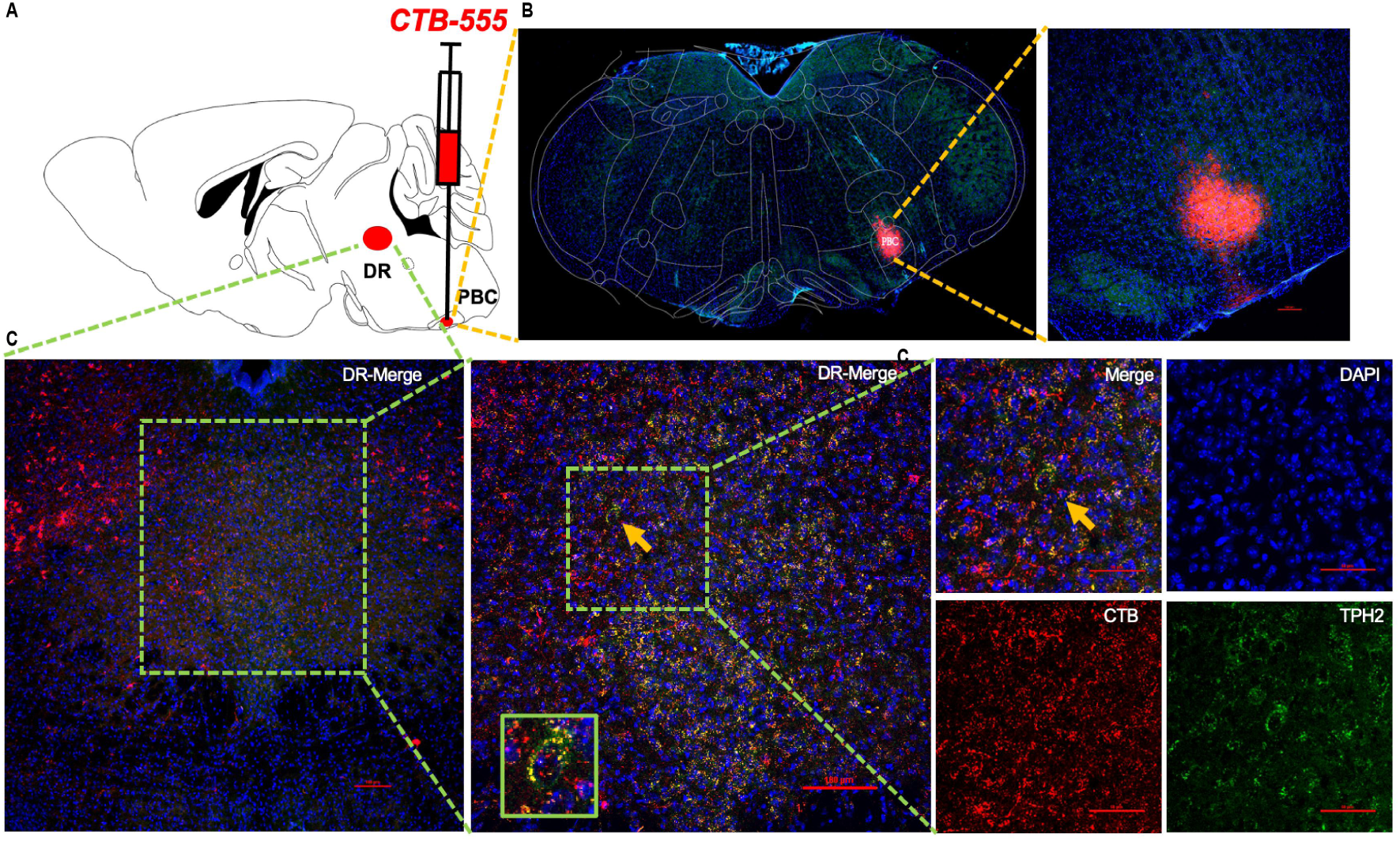
Neural projection from the PBC to the DR was established by application of the nerve tracer CTB555. A, Representative coronal brain slice, showing the location of CTB555 injection with the co-expression of NK1R and 5HT2AR in the DR. B, Projection to the DR with co-expression of TPH2 and CTB555.

### 3.7 Photostimulation increased c-fos expression in 5-HT neurons in the DR

To investigate whether photostimulation increased the excitability of 5-HT neurons in the DR, we examined the neuronal expression of c-fos, an immediate early gene that is widely accepted as a marker for neuronal activity in optogenetics studies, in the DR of DBA/1 mice with and without photostimulation. Compared with the level of c-fos expression in 5-HT neurons in the DR of DBA/1 mice without photostimulation (n=2), c-fos expression in these neurons was significantly increased with photostimulation (20-ms pulse duration, 20 Hz, at 15 mW for 20 min, n=2, *P*<0.05). This result indicated that the reduction in S-IRA by photostimulation occurs via activation of 5-HT neurons (Figure 10).

**Figure 10.**
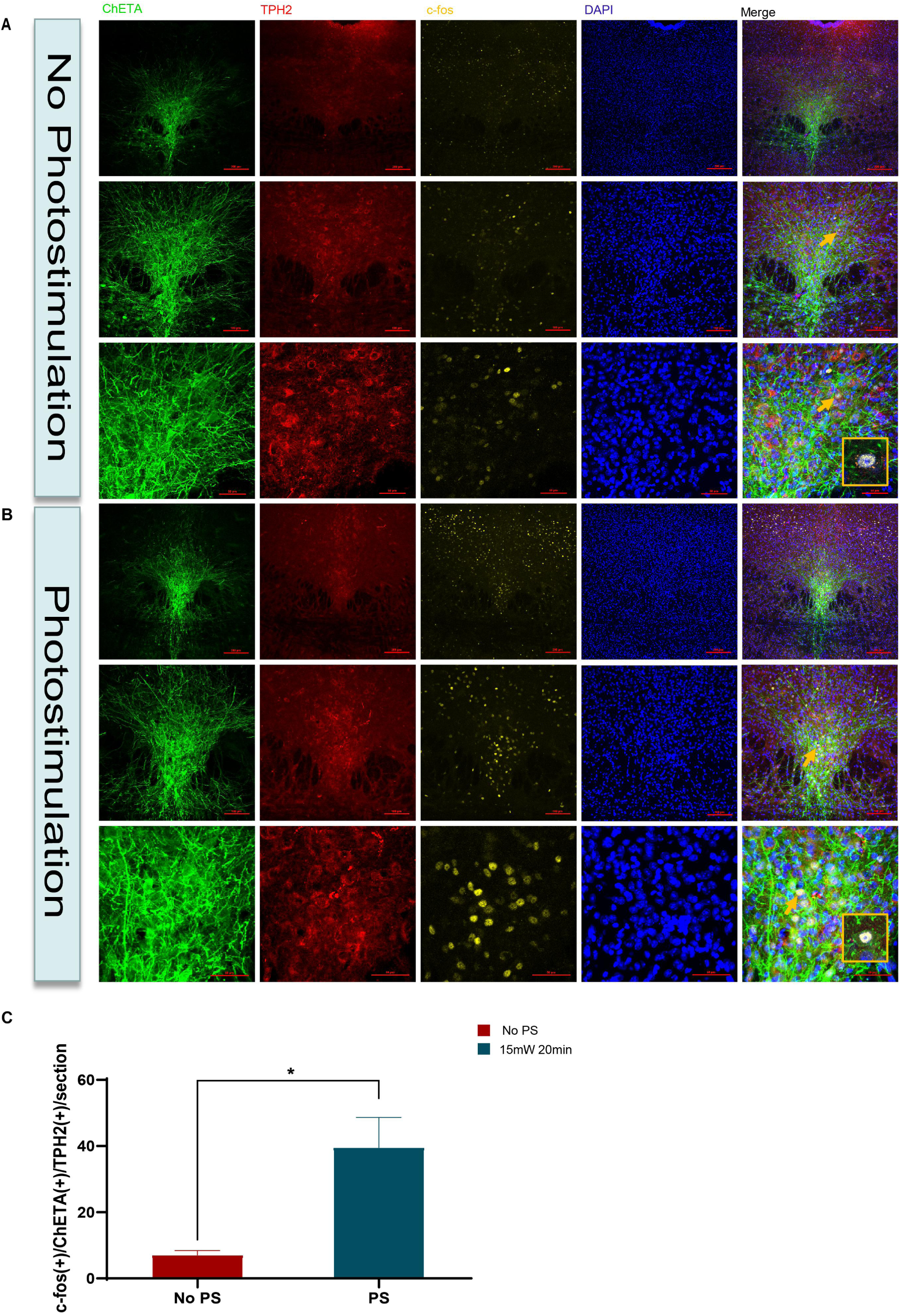
C-fos expression was significantly increased in the DR by photostimulation of TPH2-ChETA neurons in the DR of DBA/1 mice. A, Neuronal immunostaining showing co-localization of c-fos, TPH2, and GFP in a DBA/1 mouse that underwent implantation of a fiber optic cannula without photostimulation (n=2 mice). B, Immunostaining showing co-localization of c-fos, TPH2, and GFP in a DBA/1 mouse exposed to photostimulation at 15 mW for 20 min (n=2 mice). C, Quantification of c-fos(+)/GFP(+)/TPH2(+) cells in DBA/1 mice with and without photostimulation. Significantly more c-fos(+)/GFP(+)/TPH2(+) cells were observed in the PS group than in the No PS group (*P*<0.05). No PS = no photostimulation; PS = photostimulation.

### 3.8 Photostimulation of the bilateral PBC did not significantly reduce the incidence of PTZ-induced S-IRA

We next investigated whether the incidence of PTZ-induced S-IRA after photostimulation of the bilateral PBC can be independently reduced by photostimulation of TPH2-ChETA neurons in the bilateral PBC. Although the incidence of PTZ-induced S-IRA was reduced by photostimulation of the bilateral PBC, no significant difference was observed between the control group treated with PTZ and no photostimulation and the group treated with PTZ together with photostimulation (n=6 and n=7, respectively, *P*>0.05). Additionally, no significant difference was observed between the control group treated with PTZ, no photostimulation, and micro-injection of vehicle and the group treated with PTZ, photostimulation, and micro-injection of 0.4 µg KET (n=7 and n=3, respectively, *P*>0.05). These findings suggest that while activation of the neural circuit between the DR and PBC can significantly reduce the incidence of S-IRA, photostimulation of only the PBC did not significantly reduce the incidence of regulate S-IRA and SUDEP in our model (Figure 11), suggesting that the limited increment of 5-HT content caused by photostimulation of exclusively the PBC to combine 5-HT2AR in the PBC did not maintain the normal respiratory rhythm against S-IRA and SUDEP

**Figure 11.**
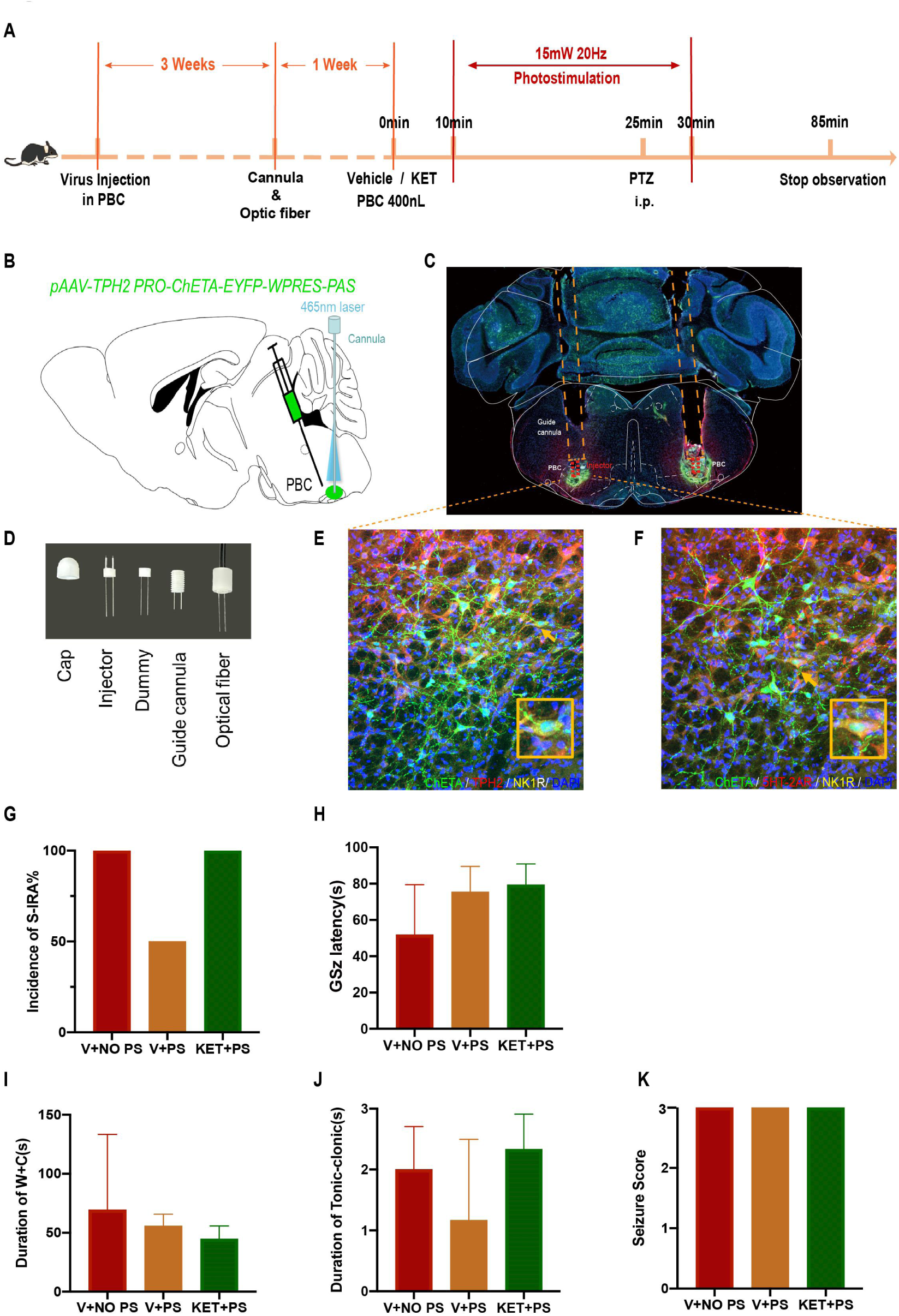
Activation of ChETA-TPH2 neurons in the PBC could suppress S-IRA but did significantly not reduce the incidence of S-IRA in DBA/1 mice or prevent death. B-F, The tracks of optic fibers with cannula implanted into each side of the bilateral PBC and staining for ChETA, TPH2, NK1R, 5-HT2AR and DAPI expression in the bilateral PBC. G, Compared with that in the control group treated with PTZ without photostimulation, the incidence of PTZ-induce S-IRA was not significantly reduced by photostimulation of the bilateral PBC (*P*>0.05). Furthermore, no significant differences were observed between the group with photostimulation and microinjection of vehicle into the bilateral PBC and the group with photostimulation and microinjection of KET into the bilateral PBC (*P*>0.05). H-K, No obvious differences between treatment groups were observed in the seizure score, duration of wild running and clonic seizure, AGSz latency, duration of tonic seizure, or seizure score (*P*>0.05). No PS = no photostimulation; PS = photostimulation.

### 3.9 PTZ-induced neuronal activity in PBC during seizures was significantly reduced by photostimulation of the DR based on photometry recordings

Calcium signaling within neurons of the bilateral PBC was recorded by photometry in mice infected with GCaMP6f in the bilateral PBC without photostimulation of the DR during the clonic and tonic seizure phases evoked by PTZ. The activity wave of calcium signals in the bilateral PBC was strong during the phases of clonic and tonic seizures in the group without photostimulation of the DR but weaker in the group with photostimulation of the DR during these phases. These data indicate that photostimulation of the DR can reduce abnormal calcium signaling activity in the bilateral PBC, which could serve to normalize respiration rhythm and thereby prevent S-IRA and SUDEP (Figure 12).

**Figure 12.**
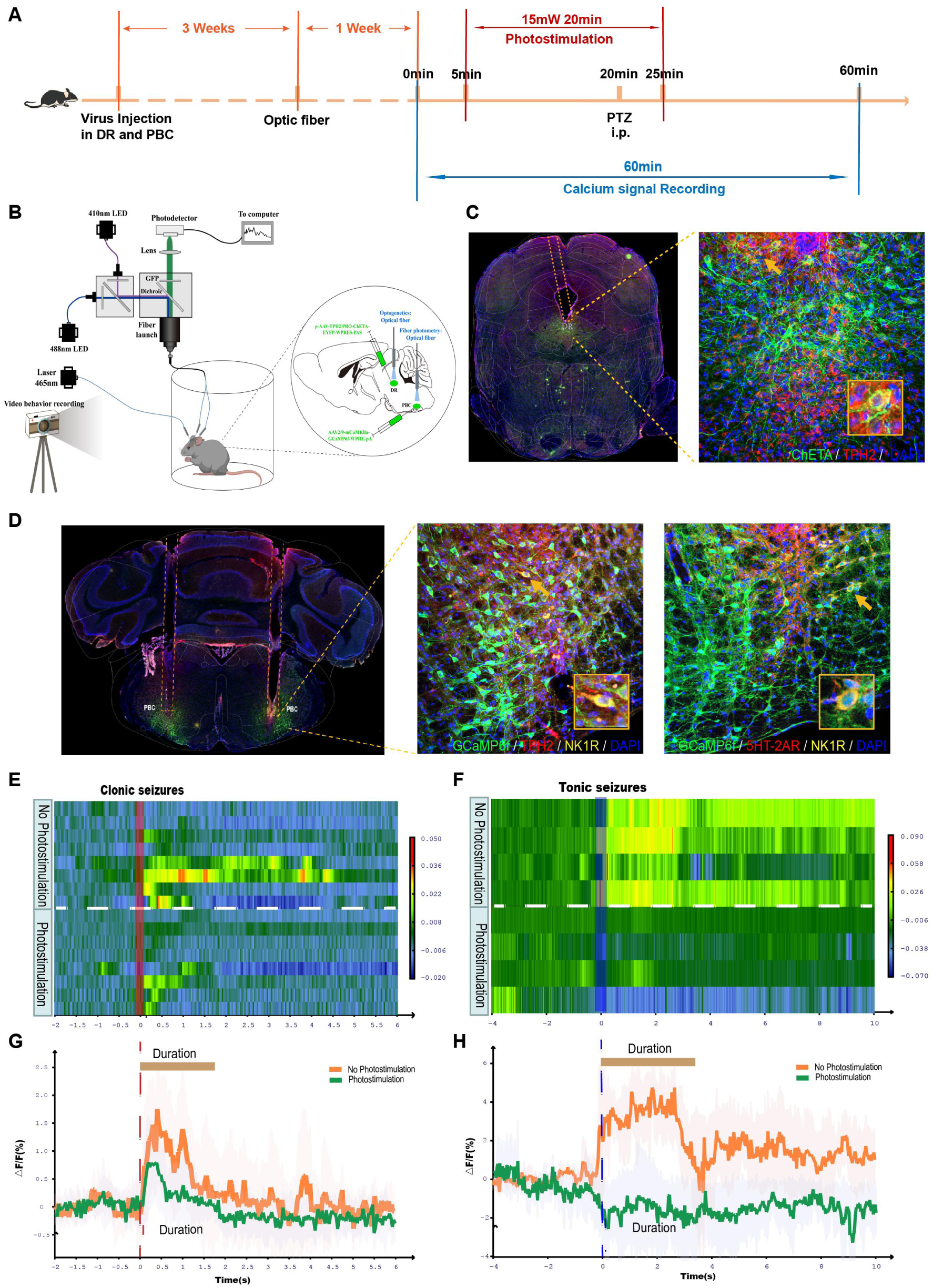
Photostimulation of the DR influenced calcium signaling in the PBC during PTZ-induced seizures. A-D, Photometric recordings from all DBA/1 mice infected with ChETA and GCaMP6f in the DR and the bilateral PBC and of neural activity based on calcium signaling in the bilateral PBC. Shown are representative images of the tracks of optic fibers implanted into the DR and the bilateral PBC and staining for ChETA, TPH2 and GCaMP6f, NK1R, 5-HT2AR, and DAPI in the DR and bilateral PBC. E-H, In the group without photostimulation of the DR, the activity wave of calcium signals from the bilateral PBC was strong during clonic and tonic seizures evoked by PTZ. E-F, However, the activity wave of calcium signals was weaker in the group with photostimulation of the DR during clonic and tonic seizures.

**Figure 13.**
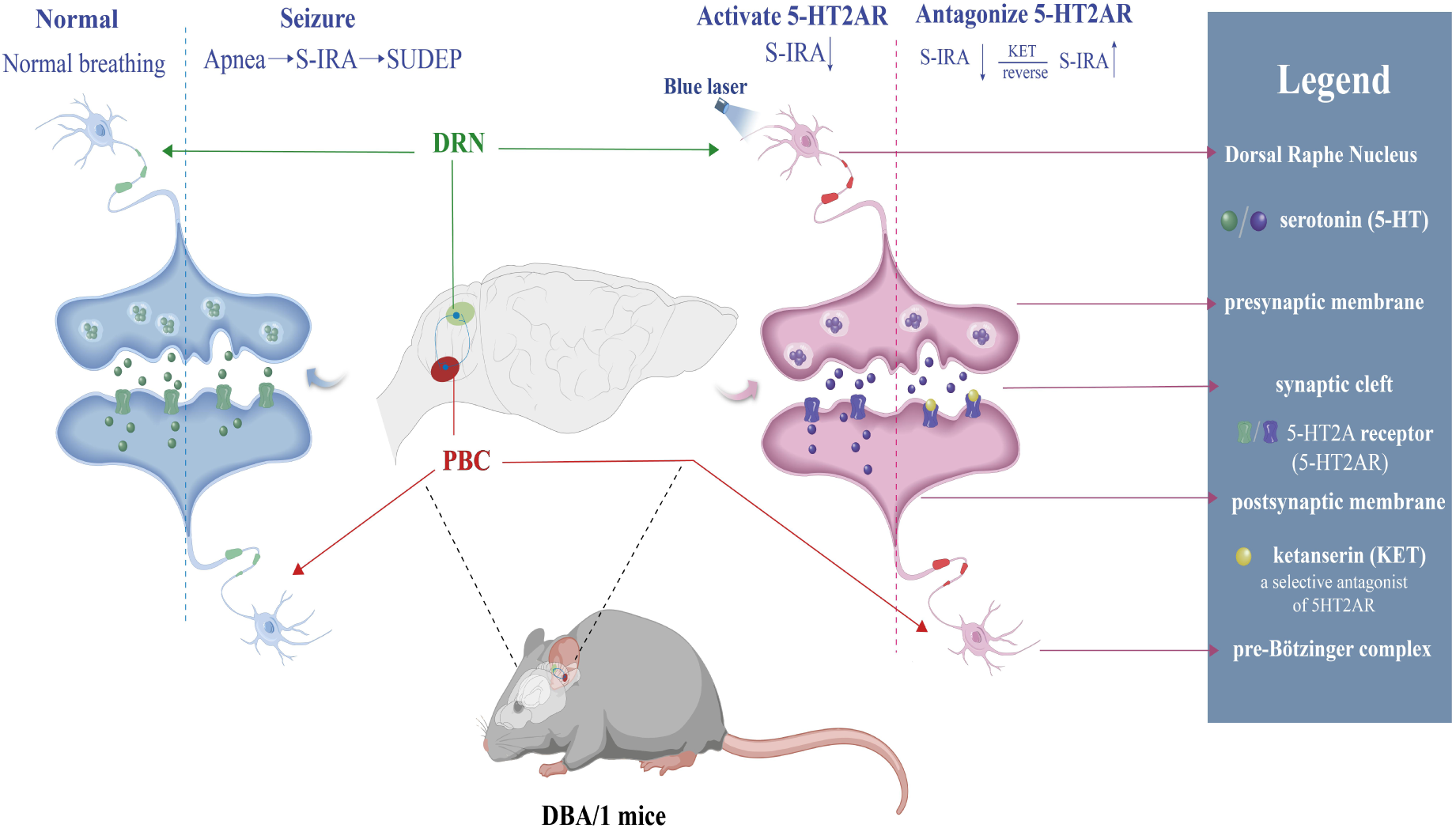
Mechanism by which 5-HT neurons in the brain modulate S-IRA and SUDEP via the neural circuit between the DR and PBC. SUDEP: sudden unexpected death in epilepsy; S-IRA; seizure-induced respiratory arrest; KET: ketanserin; PBC; pre-Bötzinger complex; DR; dorsal raphe nucleus; 5HT2AR; 5HT2A receptor.

## 4. DISCUSSION

SUDEP is a fatal complication of epilepsy, and although initial advances in understanding the role of 5-HT in the nervous system have helped to identify some causes of SUDEP, the pathogenesis of SUDEP remains poorly understood ^1–4^. Based on our previous observation that administration of 5-HTP significantly reduced the incidence of S-IRA in AGSz and PTZ SUDEP models, we next sought to further elucidate the mechanism by which 5-HT2AR in the brain may modulate the pathogenesis of S-IRA in our models. In the present study, the lower incidence of S-IRA following 5-HTP treatment was significantly reversed by IP or ICV injection of KET in our models of S-IRA induced by treatment with acoustic stimulation or PTZ, a chemoconvulsant that is widely used to model human generalized seizures. While KET reversed the suppressive effect of 5-HTP on S-IRA in a dose-dependent manner in the PTZ injection model, a ceiling effect was observed for the ability of KET to reverse the 5-HTP–mediated reduced incidence of S-IRA in the acoustic stimulation SUDEP model.

Our previous study showed that administration of 5-HTP significantly reduced the incidence of S-IRA via anti-convulsant effects ^5, 7^. This indicated that most treated DBA/1 mice avoided S-IRA without experiencing tonic–clonic seizures other than wild running and generalized clonic seizures, and thus remained sensitive to some seizures. Different from our previous study in which atomoxetine reduced the incidence of S-IRA evoked by acoustic stimulation without affecting seizure behavior in DBA/1 mice ^8–9^, administration of 5-HTP significantly reduced the incidence of S-IRA through its anticonvulsant effects in our models. Such an anticonvulsant effect of 5-HTP is basically consistent with other studies showing that activation of 5-HT neurons can reduce the severity of seizures ^16–17^. Other groups showed that boosting the 5-HT level in the brain suppresses seizures via anti-convulsant effects as well ^23–24^. However, administration of KET increased the incidence of S-IRA in response to acoustic stimulation or PTZ injection even after 5-HTP treatment with the occurrence of partly wild running, clonic and/or tonic-clonic seizures, which demonstrated that IP administration of KET reversed the reduction of the incidence of S-IRA by 5-HTP and only partly affected seizure behaviors by acting on the neuron nucleus controlling respiratory activity.

To further determine the effects of KET on the incidence of S-IRA via action on the central nucleus in the brain, we administrated KET via the ICV pathway and measured its effect on S-IRA in DBA/1 mice in both SUDEP models. The results showed that KET reversed the suppressive effects of 5-HTP both via IP injection and ICV injection, suggesting that the reversal effect of KET is independent of the SUDEP model type.

Meanwhile, most DBA/1 mice in the different treatment groups recovered from S-IRA within 24–72 h, indicating that the recovery interval for S-IRA depends on the concentration of 5-HT in the brain. In addition, we observed a dose-dependent effect on the reversal of the effect of 5-HTP on the S-IRA incidence and a ceiling effect with 25 mg/kg KET (IP). Indeed, our previous study showed that the reduced incidence of S-IRA achieved via activation of 5-HT neurons in the DR by optogenetics was significantly reversed by ondansetron, a specific antagonist of the 5-HT3 receptor ^7^. However, we cannot rule out the possibility that 5-HT2AR mediates the pathogenesis of S-IRA and SUDEP by closely interacting with 5-HT3 receptor (5-HT3R) in the brain. Further research is needed to characterize the interaction between 5-HT2AR and the 5-HT3R

Previously an SSRI also was shown to be effective at reducing S-IRA evoked by maximal electroshock (MES) in Lmx1b(f/f) mice on a primarily C57BL/6J background, a strain that is resistant to AGSz, and depletion of 5-HT neurons enhanced seizure severity, which led to S-IRA and could be prevented by 5-HT2AR activation through IP injection of with DOI (2,5-dimethoxy-4-iodophenylpropane hydrochloride), a selective agonist for 5-HT2AR, in a MES model ^18^. However, it remained unclear whether depleting 5-HT neurons itself led to S-IRA and the reversal effects by DOI targeting peripheral or central 5-HT2AR. By contrast, based on our previous findings, we further explored the role of 5-HT neurotransmission and 5-HT2AR by peripheral and central intervention approaches with KET, a selective antagonist for 5-HT2AR, in different SUDEP models. Thus, according to our findings, the role of 5-HT2AR in modulating S-IRA is supported by the results from the MES model, which further strengthens our understanding of the role of 5-HT in the pathogenesis of S-IRA and SUDEP and aid the future design of therapeutic strategies to prevent SUDEP.

Moreover, based on localized expression of 5-HT2AR in the PBC, which plays an important role in modulating respiration rhythm, and on our previous finding that the incidence of S-IRA can be significantly reduced by activation of TPH2-ChR2 neurons in the DR, we further investigated the mechanisms of how the interaction between 5-HT and 5-HT2AR mediates S-IRA and SUDEP in the same models. For this study, we used optogenetics methods to test whether activation of TPH2-ChR2 neurons in the DR significantly reduced the incidence of S-IRA evoked by PTZ via activating 5-HT2AR in the PBC. In the present study, the reduction in the incidence of PTZ-induced S-IRA via optogenetic activation of TPH2-ChETA neurons in the DR was remarkably reversed by ICV injection of KET. Subsequently, the reduction in the incidence of PTZ-induced S-IRA by photostimulation of the DR was significantly reversed by microinjection of KET into the bilateral PBC, in turn, suggesting that photostimulation of the DR remarkably reduced the incidence of S-IRA by activating 5-HT2AR in the PBC in our model. To further examine the bridge between the DR and PBC and its role in modulating S-IRA in our models, CTB-555, a nerve tracer, was used to confirm and establish the neural circuit between the DR and PBC. The results confirmed that the reduction in the incidence of PTZ-induced S-IRA upon photostimulation of the DR stems from the neural circuit between the DR and PBC. However, we only tested this using photostimulation of the DR and direct inhibition of 5-HT2AR in the bilateral PBC in this study. Whether activation of TPH2-neurons in the PBC can reduce the incidence of S-IRA still needs to be verified in subsequent experiments. Of course, the roles of other neural circuits between the DR and other brain structures involved in modulating S-IRA and SUDEP cannot be excluded in our models. Nevertheless, our pharmacologic and optogenetic findings suggest that the neural circuit between the DR and PBC plays a key role in modulating S-IRA and SUDEP.

In addition, unlike in our pharmacology experiments, although S-IRA of DBA/1 mice was blocked in optogenetic experiments, most DBA/1 mice continued to have clonic and tonic seizures. Upon analyzing the cause for this difference, the specificity of optogenetics and the specific delivery of KET by both ICV injection and injection into the PBC may be a cause. Meanwhile, the incidence of S-IRA evoked by PTZ was significantly reduced by activating the neural circuit between the DR and PBC without changing EEG activities, which further reflects the specificity of S-IRA inhibition in optogenetic experiments.

Our data also showed that activation of TPH2-ChETA neurons in the PBC is insufficient for significantly reducing the incidence of PTZ-induced S-IRA. Furthermore, the neuronal calcium signaling in the PBC during seizures induced by PTZ was significantly reduced by photostimulation of the DR according to photometry recordings. Thus, activation of the neural circuit between the DR and PBC can specifically prevent S-IRA and SUDEP.

### Conclusion

The results of the present study suggest that the lowered incidence of PTZ-induced S-IRA achieved by administration of 5-HTP can be significantly reversed by treatment with KET. Furthermore, the suppressive effects of optogenetic activation of the serotonergic neural circuit between the DR and PBC on S-IRA also were obviously reversed by KET in the DBA/1 mouse model. Therefore, 5-HT2AR in the PBC may be a specific and key target for the development of interventions to prevent SUDEP.

## 6 ACKNOWLEDGEMENTS

We are grateful to Yi Shen and Yu Dong Zhou for developing the experimental design.

